# MiniXL: An open-source, large field-of-view epifluorescence miniature microscope for mice capable of single-cell resolution and multi-brain region imaging

**DOI:** 10.1101/2024.08.16.608328

**Authors:** Pingping Zhao, Changliang Guo, Mian Xie, Liangyi Chen, Peyman Golshani, Daniel Aharoni

## Abstract

Capturing the intricate dynamics of neural activity in freely behaving animals is essential for understanding the neural mechanisms underpinning specific behaviors. Miniaturized microscopy enables investigators to track population activity at cellular level, but the field of view (FOV) of these microscopes have been limited and does not allow multiple-brain region imaging. To fill this technological gap, we have developed the eXtra Large field-of-view Miniscope (MiniXL), a 3.5g lightweight miniaturized microscope with an FOV measuring 3.5 mm in diameter and an electrically adjustable working distance of 1.9 mm ± 200 μm. We demonstrated the capability of MiniXL recording the activity of large neuronal population in both subcortical area (hippocampal dorsal CA1) and deep brain regions (medial prefrontal cortex, mPFC and nucleus accumbens, NAc). The large FOV allows simultaneous imaging of multiple brain regions such as bilateral mPFCs or mPFC and NAc during complex social behavior and tracking cells across multiple sessions. As with all microscopes in the UCLA Miniscope ecosystem, the MiniXL is fully open-source and will be shared with the neuroscience community to lower the barriers for adoption of this technology.

## Introduction

Imaging of Ca2+ indicators (*1*) with head-mounted miniature microscopes (*2–6*) has revolutionized the study of neural activity in freely moving animals, shedding light on the neural underpinnings of many behaviors such as spatial navigation (*3*), social interaction (*7*, *8*), learning and memory (*9*), sleep (*10*, *11*), and feeding (*12*). While previous iterations of UCLA Miniscopes (*13–15*), notably the V4 Miniscope (https://github.com/Aharoni-Lab/Miniscope-v4/wiki), and other open-source (*16–18*) and commercialized miniaturized microscopes have provided valuable insights by enabling the monitoring of neural dynamics with a field of view (FOV) of up to 1 mm^2^, most of them have been unable to comprehensively map the coordination of neural activity across different brain regions due to their limited FOV.

Previously, we developed an expanded FOV miniaturized microscope called the MiniLFOV (https://github.com/Aharoni-Lab/Miniscope-LFOV) with an FOV of up to 3.6 mm x 2.7mm and a spatial resolution, in none scattering media, of 2.45 μm (*14*, *19*). However, the MiniLFOV weighed 13.9 g making it suitable only for rats and larger animals. As mice are the predominant model for circuit-level neuroscience research, there is an urgent need for a lighter and more compact miniaturized microscope with a large FOV, enabling increased cell counts and multi-region imaging of neural activity stably across sessions in freely behaving mice.

In response to this need, here we introduce the MiniXL, a much lighter version of our head-mounted, open-source, large-FOV miniature microscope as part of the UCLA Miniscope Project. This new model has a 3.5 mm diameter FOV, with a 4.4-μm lateral resolution, weighs only 3.5 grams and is only 30 mm in height. Its front working distance (WD) is electrically adjustable, with a range of 1.9 mm ± 200 μm, facilitated by an electrowetting lens (EWL). Additionally, the MiniXL integrates a nine-axis absolute head-orientation sensor, capable of capturing head movement data at up to 100 Hz, which is instrumental in exploring head orientation–related neural mechanisms.

We tested the MiniXL in freely behaving mice. We imaged GCaMP6f-expressing neurons in the dorsal hippocampal CA1 pyramidal layer in both one-dimensional and two-dimensional environments, demonstrating robust spatial coding across large population of neurons. Excitingly, we simultaneously measured neural activity in left and right medial prefrontal cortex (mPFC) during a free social interaction task which could be conducted across multiple sessions and days. In addition, we simultaneously imaged neurons in the mPFC and nucleus accumbens (NAc). These experiments confirmed the MiniXL’s stability for long-term cell tracking and capability for multi-region imaging during complex behaviors.

## Results

### System design and optical performance

The MiniXL platform is comprised of an optics module and a custom rigid-flex printed circuit board (PCB). The PCB integrates several crucial subcircuits: a light-emitting diode (LED) and constant current LED driver for excitation light delivery, electrowetting lens (EWL) and EWL driver for focus adjustment, absolute head-orientation sensor, power distributor, and a complementary metal oxide semiconductor (CMOS) image sensor for signal capture (Fig.1A). This assembly is coupled to an optics module containing a quartet of achromatic lenses (#63692, #63691, #45262 × 2, Edmund Optics), in line with an excitation filter (ET470/40x, Chroma), a dichroic mirror (T495lpxr, Chroma), an emission filter (ET525/50m, Chroma), and a 3D-printed lens holder incorporating an EWL driver (MAX14515, Maxim) for precise focus control. Leveraging a 5 megapixels (MP) monochrome CMOS sensor (MT9P031, Onsemi), the system captures fluorescence signals across a substantial field of view, facilitated by power-over-coax technology and a sophisticated serializer/deserializer pair (DS90UB913A/DS90UB914A, Texas Instruments) based on a single, flexible 50-ohm coaxial cable (CW2040-3650SR, CoonerWire). This arrangement not only ensures streamlined power management and bidirectional communication but also enables high-bandwidth, unidirectional data streaming essential for capturing dynamic neural processes. The system’s compatibility with the UCLA Miniscope Data Acquisition (DAQ) hardware and software extends its functionality, allowing for backwards-compatible real-time video streaming and recording, and also enables multi-stream neural and behavioral recording. This DAQ platform supports adjustment of imaging parameters (excitation intensity, EWL focus, image sensor gain, and frame rate), real-time ΔF/F display, real-time pose estimation (*20*), and synchronization with external devices, enhancing experimental versatility.

**Fig. 1.**
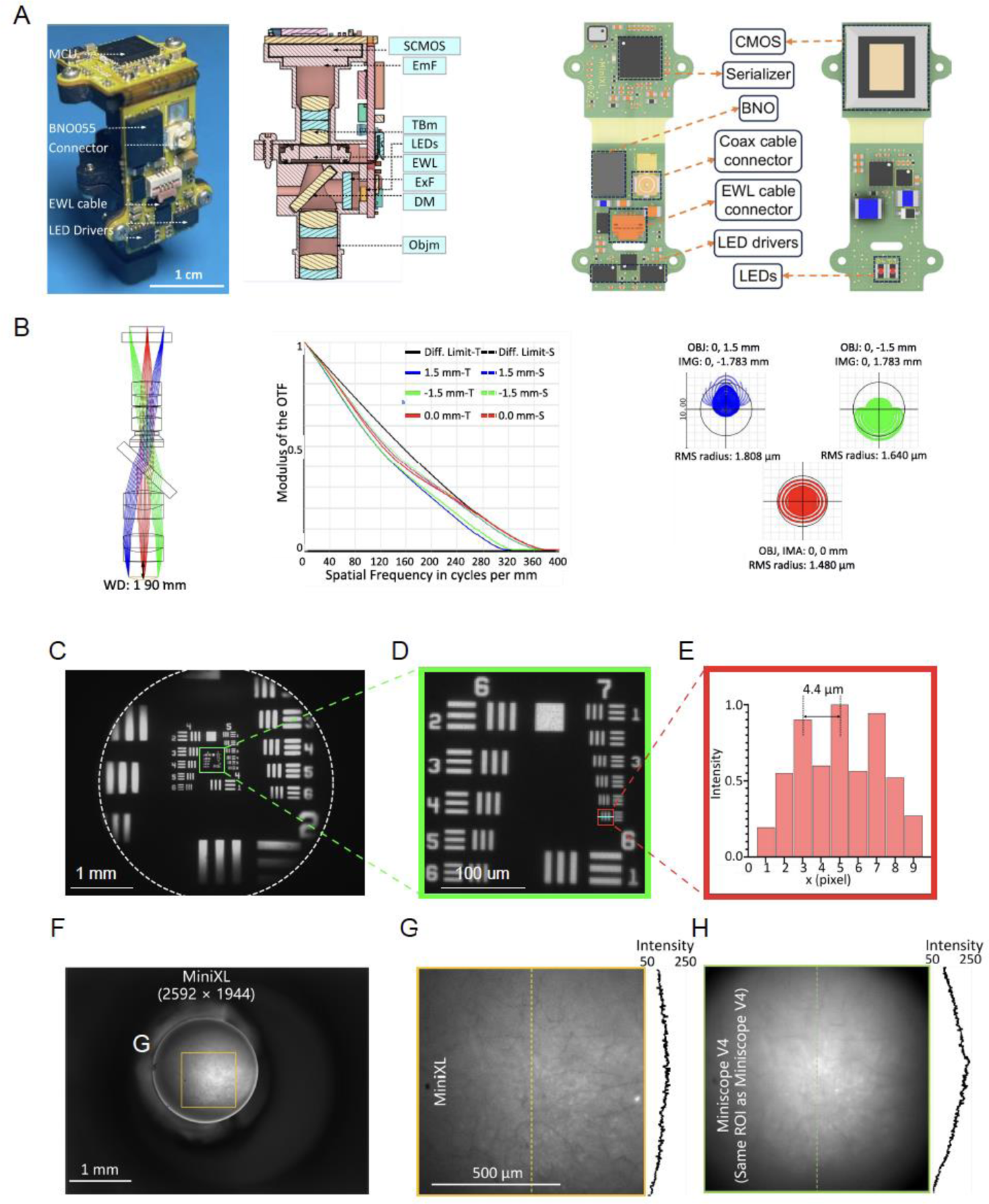
Optical design and performance of MiniXL. (A) Left panel: photograph of MiniXL. Middle panel: Cross section profile. Right panel: PCB layout. Scale bar, 1 cm. (B) Zemax simulation of emission path shows a 1.9 mm working distance (WD) with 300-μm field curvature across a 3-mm diameter FOV in the object space (left). MTFs on the image plane (top right), and spot diagram of the optics (left panel), MTFs on the image plane (middle), and spot diagram of the optics (right). In the spot diagram, root mean square (RMS) radius is 1.480 μm at the center and 1.808/1.640 μm at (0, 1.5 mm/−1.5 mm). The magnification of the optics is given by 1.19 calculated from the spot diagram. (C-E) FOV (3.5 mm in diameter) shown using the Ø1” 1951 USAF Targe (#R1DS1N, Thorlabs). Scale bars, 1 mm (C) and 100 μm (D). The value 4.4 μm (group 7 element 6, 228 lps/mm) can be resolved from the green line in (D) (E). (F-H) Comparison of MiniXL (F, G) and Miniscope V4 (H) in the same ROI. The field of view of Miniscope V4 is 1.1 mm in diameter. MiniXL shows more uniform fluorescent detection than Miniscope V4.

Additionally, the system’s optical design (Fig. 1A), is designed to achieve an optical resolution up to 2.5 μm across a 3 mm diameter FOV by utilizing off-the-shelf lenses. The MiniXL incorporates a Rigid-flex PCB design, comprising four rigid PCBs interconnected by an internal flex-printed circuit. The LED circuit board accommodates two LEDs, which are powered by dedicated LED drivers featuring an I2C digital potentiometer for precise excitation light adjustment. Adjacent to the LED assembly, an EWL driver and an EWL cable secure the EWL device, alongside an absolute orientation sensor (BNO055, Bosch Sensortec) for real-time collection of head orientation data. Additionally, a high-resolution 5-MP monochromatic CMOS image sensor (MT9P031, onsemi) is employed to capture Ca2+ fluorescence, transmitting digitized image data to the serializer system. This system further processes and serializes the imaging data, transmitting it via a coaxial connector for seamless integration with a custom Miniscope DAQ system. Its design aims to achieve a balance between resolution, sensitivity, and field coverage while facilitating unconstrained animal behavior, enabling an electrically adjustable working distance (1.9 mm ± 200 μm) to accommodate different imaging depths (Fig. 1B). At its core, the system’s 2.2-μm pixel-size, 5 MP CMOS image sensor expands the achievable field of view while maintaining small form factor optics and cellular resolution (Fig. 1C, D, E). Additionally, the MiniXL has significantly improved imaging FOV, optical contrast and even detection in fluorescence than that of a V4 Miniscope (Fig. 1F-H).

The MiniXL weighs 3.5 g, similar in weight to many other existing head-mounted neural recording devices for freely behaving mice (*2*, *13*, *15*, *18*), and does not hinder free behavior (Fig. S1). Below, we show the MiniXL successfully used in freely behaving animals across a number of behavioral tasks, including open field test, linear track, and social interactions paradigms.

### Imaging dorsal CA1 (dCA1) place cells in freely moving mice in both 1D and 2D environments across days

To evaluate the MiniXL’s capabilities, we first imaged GCaMP6f-labeled neurons within the dCA1 region, known for its critical role in spatial navigation. Recording place cell activity in CA1 allows us to compare the performance of this microscope with previous versions of miniature microscopes, specifically the UCLA Miniscope versions V3 and V4 and MiniLFOV (*3*, *13*, *14*). We leveraged the large FOV provided by the MiniXL, coupled with implantation of a stacked pair of 1.8 mm diameter GRIN lenses (Edmund optics, quarter pitch, #64-531) resulting in a relay GRIN lens configuration (Fig. 2A). These 1.8 mm or 2 mm large size GRIN lenses have been used frequently for CA1 imaging (*13*, *21*). The incorporation of a large lens and FOV enabled the acquisition of large-scale neural dynamics from 1640 neurons across three sessions (Fig. 2C-E, Movie S1).

**Figure 2.**
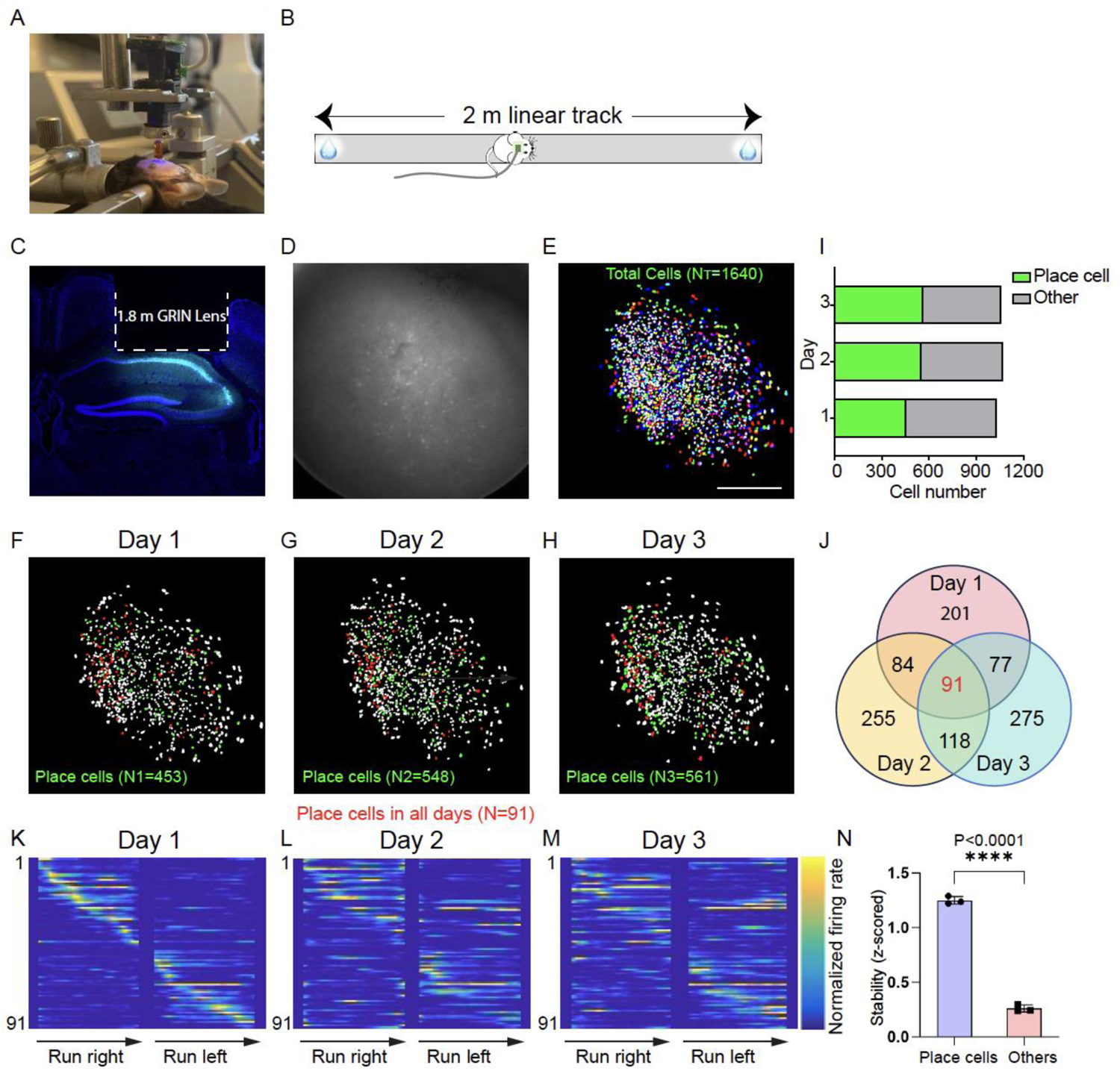
Imaging the activity of dCA1 place cells on a linear track. (A) Surgical implantation of GRIN lenses and baseplate. Two GRIN lens were stacked with the bottom one implanted upon dCA1 region. (B) Animals were trained to run on a 2-m linear track for water reward on two ends. (C) Example brain slice showing GCaMp6f expression and GRIN lens implantation track. (D, E) Example raw image of field of view and all cells identified across three days (N=1640). Each field of view is cropped to approximately 1.393 mm × 1.393 mm. (F-H) Identified place cells (green) on Day1 (N1=453), Day2 (N=548) and Day3 (N=561). There are 91 cells were identified as place cells and were matched across all sessions (red). (I) Ratio of place cells and the other cells on each day. (J) Overlap of place cells between every two days and among three days. (K-M) Normalized neural activity rates of place cells on Day 1 (K) that were also active on Day2 (L) and Day 3 (M). cells were sorted by the peak firing rate from Day 1. (N) Mean stability of place cells and other cells on each day of one example mouse. Place cells showed significant higher stability than other cells (unpaired Student’s t-test, P <0.0001).

Through this setup, we were able to capture the activity of GCaMP6f-expressing neurons in dCA1 as mice navigated a 2-meter-long linear track (Fig. 2B-D). The imaging was conducted at a rate of 23 frames per second (fps) over 15-minute intervals, generating high-resolution videos (1000 pixels x 1000 pixels for each frame) that were then cropped to the GRIN lens’s diameter size (1.8-mm in diameter, 704 pixels x 704 pixels) for analysis. Concurrently, a synchronized behavioral camera recorded the mouse’s position at 50 fps, providing a synchronized neural-behavioral dataset for analysis. We used CalmAn (https://github.com/flatironinstitute/CaImAn) and CellReg (*22*), analysis packages which identified, demixed, deconvolved, and tracked 1640 neurons in total across three sessions on different days. A significant proportion of neurons recorded demonstrated spatial tuning in the linear track (44.02% on Day 1; 51.21% on Day 2; 53.07% on Day 3) (Fig. 2F-J). The percentage of place cells we have identified is comparable to that reported in previous studies using miniscope calcium imaging (*13*). This dataset allowed us to identify and track place cells across sessions, showcasing their stability and spatial encoding properties (Fig. 2K-N).

We also tested the MiniXL by recording CA1 neural dynamics during exploration of a 2-dimensional (2D) open-field arena (Fig 3A, B). Across a 30-minute recording session, the MiniXL data yielded 764 cells (Fig. 3C, D), with more than one third of neurons ((N =266; 34.82%) meeting the criteria (see **Methods**) for place cell identification (Fig. 3D). The analysis highlighted clear spatial preferences and stable activity patterns among identified place cells (Fig. 3E, F), underscoring the MiniXL’s utility in exploring complex neural phenomena across diverse environment contexts. Additionally, the combined open field place field coverage from all 266 cells fully covered the arena area (Fig. 3G). Mouse speed and time spent in the center of the arena were similar between animals wearing and not wearing a MiniXL (Fig. S1A-C), indicating that MiniXL does not significantly affect locomotion or alter anxiety-related behaviors. This comprehensive approach underscores the MiniXL’s potential in advancing our understanding of neural circuitry and behavior.

**Figure. 3.**
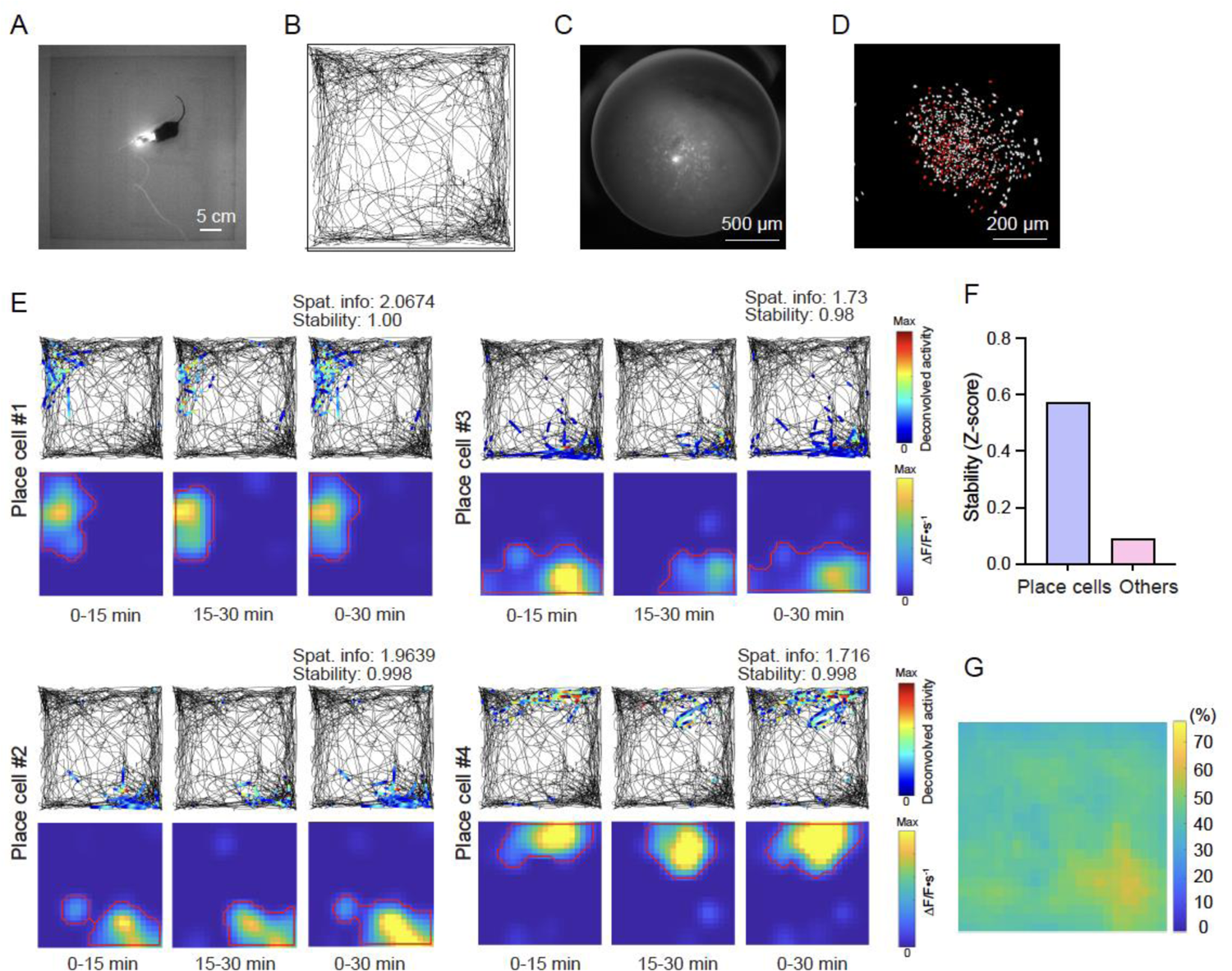
Imaging place cells in the open field arena. (A) Snapshot of mouse behavioral recording in open field arena. (B) The trajectory of mouse running in the arena. (C) Raw calcium fluorescence of the field of view from example animal. (D) Single neurons extracted from the field of view by CNMF-E with identified place cells indicated in red. (E) Example of four places cells with spatial information value and stability score. (F) Comparison of the stability score of place cells and other cells. (G) Environment coverage of place fields.

### Bilateral imaging of mPFC in freely behaving socially interacting mice

The MiniXL’s large field of view supports simultaneous imaging of more than one relay lens when lenses are located within 3.5 mm distance. Leveraging this ability, we recorded neural activity from bilateral mPFCs during mouse social interactions (Fig. 4C). We implanted two 1 mm diameter, 4 mm long relay lenses (Inscopix, 1050-004595) above the right and left prelimbic area of the mPFC labeled with GCaMP6f (Fig. 4A, B, E, F, Movie. S2), and performed calcium imaging as mice engaged in social or object interactions. As expected, mice engaged more frequently in social interactions (23.7% of total time) compared to object exploration (2.4% of total time) (Fig. 4C, D). This result is comparable to results obtained in mice implanted with a single relay lens in the mPFC or in mice not implanted with any relay lenses (Fig. S2). Wearing a MiniXL did not significantly affect mouse behavior, including the time spent interacting with social target or the percentage of time spent sniffing, running, rearing, self-grooming or resting (Fig. S1E-G). Neuronal activity, recorded during these interactions and counterbalanced with object exploration sessions, revealed that a proportion of neurons were significantly activated or inhibited by social or object interaction (Fig. 4G-J) (Left: socially excited neurons (SE), 11.99±0.02%; socially inhibited neurons (SI), 7.92±0.02%; object excited neurons (OE), 4.82±0.01%; object inhibited neurons (OI) 3.60±0.01%; Right: SE, 11.56±0.01%; SI, 5.96±0.02%; OE, 6.22±0.02%; OI, 6.35±0.02%). Our findings suggest a similar distribution of neurons responsive to social interactions and object exploration across both hemispheres (Fig. 4K). Notably, the inter-hemispheric neural activity correlation (Pearson Correlation Coefficient) in the mPFC was higher during social interactions than in object exploration sessions (Fig, 4L, All cells). Higher correlations were particularly evident among neurons excited or inhibited during social engagement (Fig, 4L, Excited cells, Inhibited cells) and there was no significant difference during non-interaction period between social session and object session (Fig. S3) though no lateralization in social encoding was observed. This underscores a more synchronized neural response in the PFC during social behaviors, highlighting the complex neural coordination underlying social interactions. Moreover, we could track the same cells across multiple days during mice freely social interactions, attesting to the stability of MiniXL imaging (Fig. S4).

**Figure. 4.**
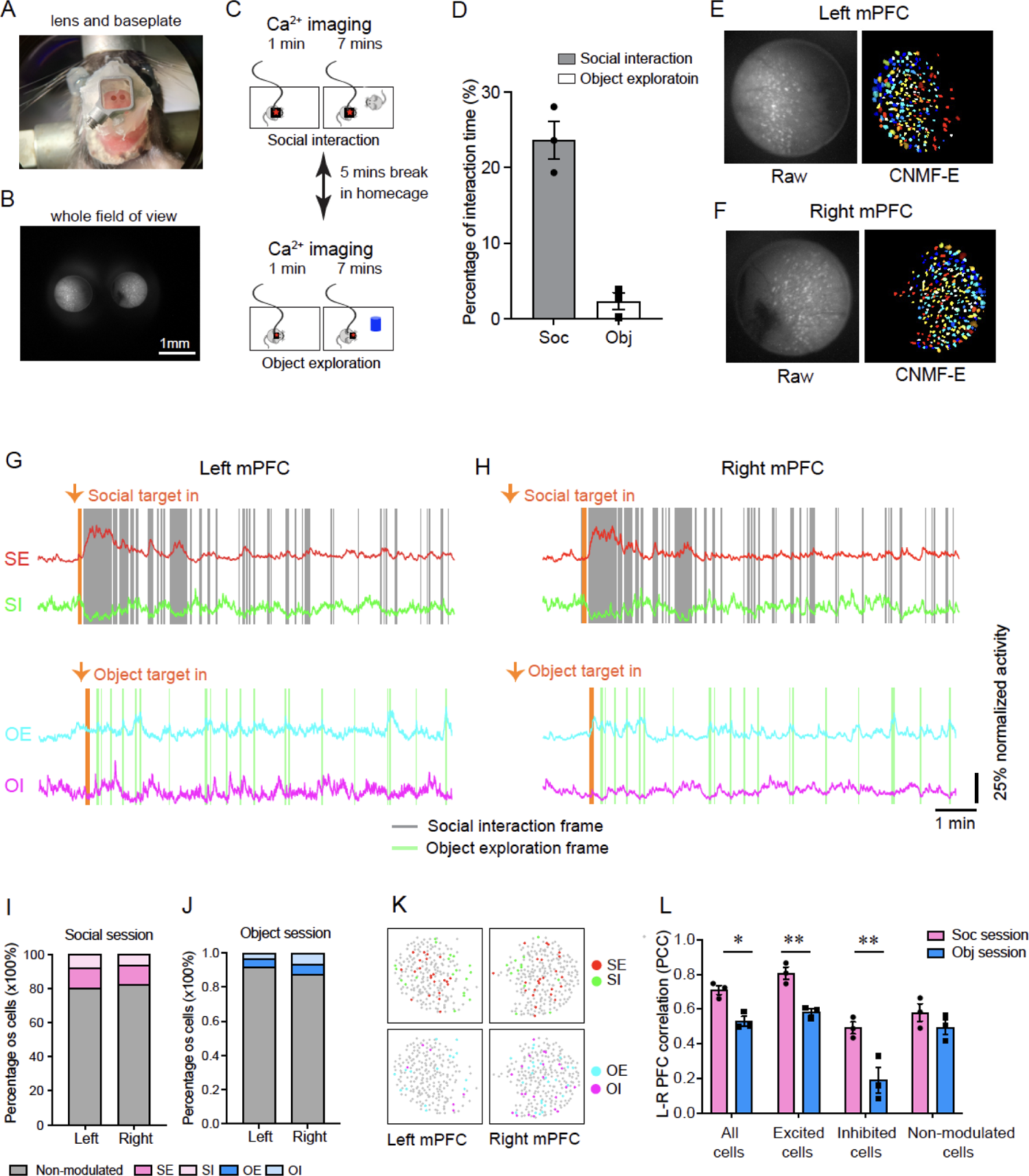
Bilateral imaging of PFC in freely behaving mice using MiniXL. (A) Diagram of bilateral lens implantation above left and right mPFC and baseplate installation. (B) MiniXL’s FOV across dual GRIN relay lens implants above mPFC. (C) Schematic diagram of social interaction session counterbalanced with object exploration during calcium imaging with MiniXL. (D) Percentage of time that animal spent on social interaction and object exploration. (E, F) Raw calcium fluorescence frame from an example animal and single neurons extracted from the FOV. (G, H) Mean calcium traces of social excited cells (SE), social inhibited cells (SI), object excited cells (OE) and object inhibited cells (OI) recorded from example mouse’s left and right mPFC. (I, J) Percentage of SE, SI, OE, OI and social/object non-modulated cells. (K) Distribution of SE, SI, OE and OI in left and right mPFC. (L) Comparison of the correlation of left and right mPFC during social interaction session and object exploration session.

### Simultaneous imaging of mPFC and NAc core with the MiniXL

Understanding the neural dynamics driving behavior requires measuring coordinated activity patterns across multiple brain regions within a network. To highlight this capability, we performed simultaneous imaging of mPFC and nucleus accumbens (NAc) core during an open field test (Fig. 5), demonstrating the technical feasibility of dual-region imaging. Neurons within both the mPFC and NAc core were labeled with GCaMP6f. This was achieved through the careful choice of relay lenses: a 0.5mm diameter, 6.1mm long lens (Inscopix, 1050-004599) for the mPFC, and a 0.5mm diameter, 8.4mm long lens (Inscopix, 1050-004600) for the NAc, ensuring comprehensive coverage of the targeted areas. (Fig. 5A, B, D, Movie. S3).

**Fig 5.**
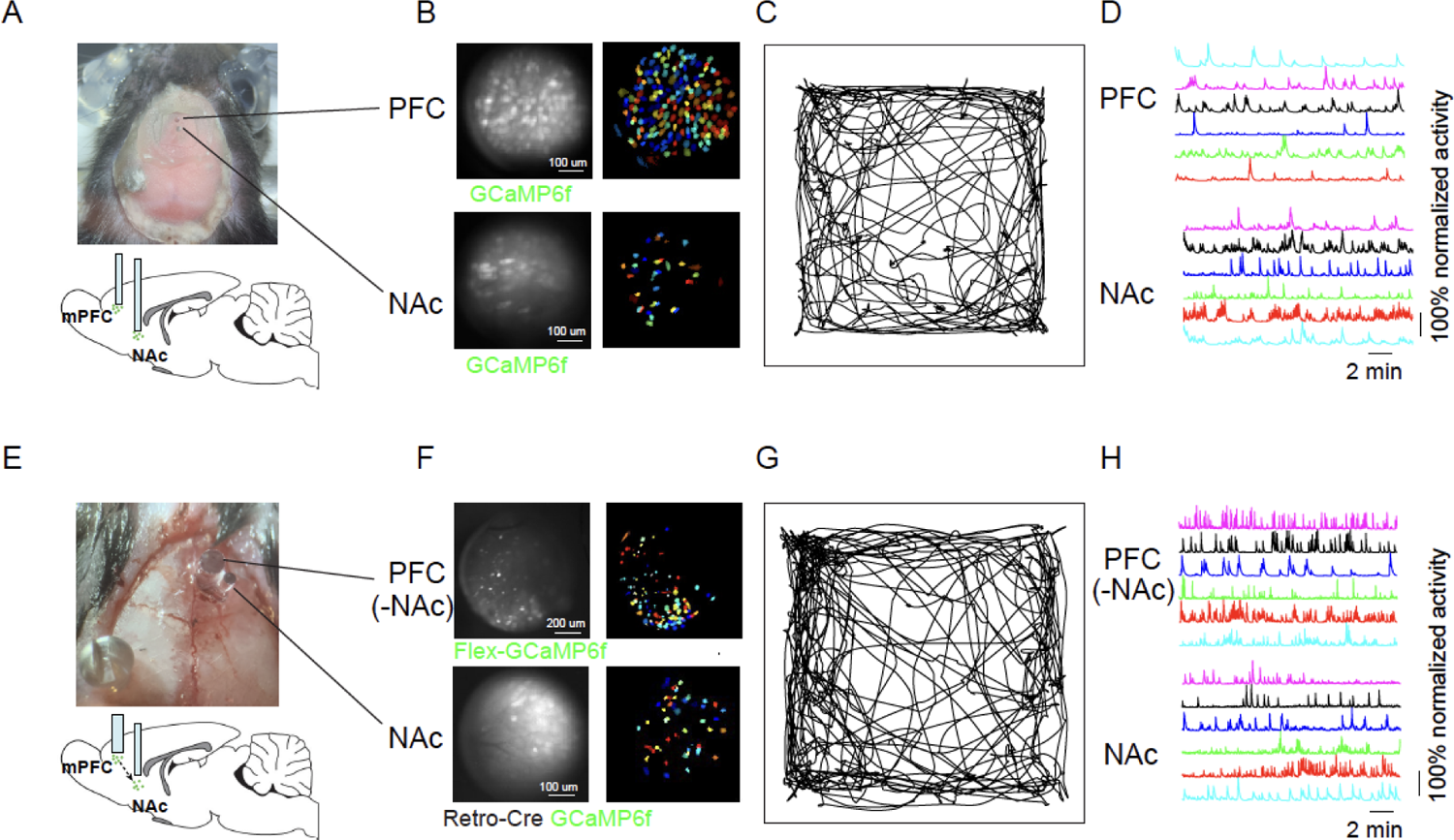
PFC-NAc simultaneous imaging by MiniXL. (A, E) Diagram of lens implantation in PFC (0.5mm/6.1mm in A and 1mm/4mm in E) and Nac (0.5mm/8.4mm). A, NAc: GCaMP6f, PFC: GCaMP6f; E, NAc: retro-Cre+GCaMP6f, PFC: Flex-GCaMP6f. (B, F) Raw calcium fluorescence frame of the FOV from example animals and single neurons extracted from the FOV. (C, G) Trajectory of mouse in open field test. (D, H) Example of calcium traces recorded from NAc and mPFC.

Given the known direct projections between the PFC and NAc, which encodes a blend of social and spatial information (*23*), we adopted a specific genetic labeling strategy. This involved the injection of a retrograde AAV expressing Cre recombinase into NAc, combined with injections of AAV-Syn-Flex-GCaMP6f in the mPFC (Fig. 5E-H). Because of the topological distribution of mPFC neurons projecting to different downstream target, we implanted 1mm relay lens above mPFC to increase the chance of recording from target cells. This approach selectively labels mPFC neurons projecting to the NAc, illuminating the upstream-downstream dynamics within this circuit. Utilizing the MiniXL, we successfully imaged the activity of mPFC neurons projecting to NAc simultaneously with the activity of NAc neurons themselves.

Combined with different virus strategies, this dual-region imaging technique could become a versatile tool for exploring neural circuits of various behavior and enrich our understanding of underlying neural mechanisms.

## Discussion

The MiniXL marks a significant advance in miniaturized microscopy imaging technology, offering several key enhancements over existing miniature microscope platforms such as the UCLA Miniscope versions V3 and V4, as well as the TINIscope (*24*), the Kiloscope (*17*) and CM^2^ (*25*). Its design enables imaging from a large FOV (3.5 mm diameter), at cellular resolution (4.4 µm) at 23 fps while maintaining a low weight (3.5 g). The adjustable working distance of 1.9 mm ±200 µm, and a novel absolute head-orientation sensor further increase the flexibility and capability of the microscope. The previous released Kiloscope also provides cellular resolution imaging across a large FOV and an even lighter weight. However, the aspheric lens assembly (originally used for smartphone) and the optical fiber results in difficulty assembly and less flexible tethering during behavior. All parts for a MiniXL are off-the shelf products and are easy to assemble. The MiniXL’s tether, a single coaxial cable with 0.3 mm in diameter, supports a wider range of natural behavior and also makes it backwards compatible with all UCLA Miniscope DAQs. This innovative platform facilitates diverse imaging modalities, enabling both deep and superficial brain region long-term imaging.

Through in vivo testing, we have validated the MiniXL’s capability to capture neuronal activity across various brain regions, including the dorsal CA1 of the hippocampus, bilateral medial prefrontal cortex (mPFC), and mPFC and nucleus accumbens (NAc) simultaneously. Complementary to previous studies showing persistent imaging of a large number of bilateral hippocampal neurons (*26*), a single MiniXL has the ability to simultaneous record more than 500 neurons from unilateral CA or nearly 500 neurons from bilateral mPFC in freely behaving mice within a single session, significantly expanding the population recorded and allowing for more comprehensive studies of neural population dynamics across multiple brain regions.

Furthermore, the MiniXL, supported by an array of online training resources and documentation provided by the UCLA Miniscope project (https://github.com/Aharoni-Lab), presents an open-source, user-friendly, and cost-effective solution for the neuroscience community. This accessibility encourages the broader adoption of calcium imaging techniques in research laboratories, fostering further discoveries in neural behavior and function while minimizing technical and economic barriers.

The MiniXL also sets a new standard for multi-region brain imaging, essential for understanding the complex interactions within neural circuits that underpin behavior. Its dual-region imaging capability, highlighted here by simultaneous imaging of the mPFC and NAc, addresses a critical need for tools that can explore the neural basis of behaviors coordinated by multiple brain regions. This feature is particularly relevant for studies on social behavior, learning and memory, and so on, where understanding the interplay between different areas of the brain and the information flow in the network is the key to unraveling the mechanisms of social interaction.

In comparison to other devices like the TINIscope and Kiloscope, the MiniXL stands out for its single-coaxial-cable tethering and electronic focal plane adjustment, which simplifies the experimental setup and minimizes interference with natural animal behaviors. Additionally, the MiniXL is backwards compatible with the UCLA Miniscope DAQ hardware and software as well as can be adopted to use the UCLA Miniscope’s optional wirefree expansion module for rats and larger animals (*14*). Its high frame rate and large FOV make it an excellent fit for studies requiring detailed imaging of complex neural networks. UCLA Miniscope project has released supportive information as usual to assist users in adopting the technology. (Github: https://github.com/Aharoni-Lab/MiniXL). In conclusion, the MiniXL represents a significant advance in in vivo neural imaging technology.

## Methods

### MiniXL design, manufacturing, and assembly

MiniXL was designed using Zemax Optics Studio. The Numerical Aperture (NA) was set to be 0.12 in the objective space and FOV was set at a minimum of 3 mm diameter. The distance between chosen lenses were optimized to achieve <4 µm resolution across at least 3-mm FOV, while maintaining the weight of scope <4 g for mice. Two achromatic lenses (#63-692, #63-691, Edmund Optics) were used to build the 1.9-mm-WD objective module. Two achromatic lens (#45-262, Edmund Optics) were adopted to build the emission module. The EWL (Corning Arctic 25N, Varioptics/Corning) is placed between the objective and emission modules for electronic focus adjustment. The MiniXL bodies including objective module, emission module, filter cover, baseplate, were designed in Autodesk Fusion 360 (educational license) and printed with black resin (FLGPBK04, Formlabs) by a stereolithographic 3D printer (Form 3, Formlabs).

Three filter slots for an excitation filter (4×4×1.1 mm, ET470/40x, Chroma), a dichroic filter (4 ×6×1mm, T495lpxr, Chroma), and an emission filter (5 ×5×1mm, ET525/50m, Chroma) were designed on the microscope bodies. All the optical components could be easily assembled without the need for any epoxy or optical glue. The circuit schematic and rigid-flex PCB layout were designed using KiCad, a free software suite for electronic design automation. The PCB was divided up into two rigid subcircuits. One was an excitation circuit including LEDs, an EWL tuning and head orientation chip. The other was a CMOS image sensor circuit with a serializer chip. The two subcircuits were connected by a double-sided embedded flex printed circuit. The assembly of all the components uses seven M1 thread-forming screws (96817a704, McMaster-Carr).

### Animals

All experimental protocols were approved by the Chancellor’s Animal Research Committee of the University of California, Los Angeles, in accordance with the NIH guidelines. All mice used here were male C57BL6/J mice obtained from Jackson Laboratory (strain 000664). Mice were older than 6 weeks old. They were maintained on a 12h:12h light/dark cycle with food and water ad libitum (except the mice doing linear track). Mice were single housed after surgery for three to four weeks before in vivo calcium imaging and behavior experiments.

### Stereotaxic surgeries

Mice were anaesthetized with 1 to 2% isoflurane-oxygen mixture and placed into a stereotactic frame (David Kopf Instruments). AAV1.Syn.GCaMP6f.WPRE.SV40 was unilaterally injected into dCA1 (400nl) (AP-2.1mm, ML 2mm, DV −1.65mm), bilaterally injected into mPFC (prelimbic cortex, PL) (300nl each) (AP +1.8mm, ML 0.4mm, DV - 2.1mm) or injected into mPFC (400nl) (AP +1.9mm, ML 0.5mm, DV - 2.5mm) and NAc (400nl) (AP +1.3mm, ML 1mm, DV - 4.7mm) of six- to seven-week-old mice at 60nl min^-1^using a Nanoject microinjector (Drummond Scientific). Alternatively, retro-Cre (400nl) and Syn.GCaMP6f (350nl) were injected into NAc and Flex-GCaMP6f (400nl) was injected into mPFC. Five to seven days after virus injection, mice were implanted with a 1.8mm diameter, 4.7mm length GRIN lens (Edmund Optics) over dCA1(coordinates of lens center: AP - 2.1mm, ML 1.8mm, DV - 1.25mm) or 1mm diameter 4mm length relay lens (Inscopix) over mPFC in each hemisphere (coordinates of lens center: mPFC: AP+1.8mm, ML 0.5mm, DV - 1.8mm); 0.5mm diameter, 6.1mm length relay lens in mPFC and 0.5mm diameter, 8.4mm length relay lens in NAc (coordinates of lens center: mPFC: AP +1.9mm, ML +0.5mm, DV - 2.5mm; NAc: AP +1.3mm, ML 0.75mm, DV - 4.3mm); 1mm diameter, 4mm length relay lens in mPFC and 0.5mm diameter, 6.1mm length relay lens in NAc (coordinates of lens center: PFC: AP +1.9mm, ML +0.5mm, D - 2.5mm; NAc: AP +1.3mm, ML +1mm, DV - 4.7mm). Lenses were secured to the skull using cyanoacrylate glue and dental cement and covered with Kwik-Sil (WPI). Two to three weeks later, MiniXL was attached to an aluminum baseplate and placed on top of the GRIN lens. After adjusting the focal plane, we secured the baseplate with dental cement. Lenses were protected by plastic cup over baseplate.

### Behavioral tests Open field test

Mice with GRIN lenses implanted in dCA1 or mPFC and NAc were habituated to the environment and to wearing MiniXL for six days before test. On Day 1 and Day 2, mice wore dummy scopes weighing 3.5 g in the homecage for one hour. On Day 3 to Day 6, mice wore dummy scopes in the homecage for 30 mins and in an open arena (45cm X 45cm X 30cm) with visual cues on the walls for 30mins. On the day of test, the microscope was attached to the baseplate and we recorded calcium signals and behavioral videos simultaneously when mice are freely moving in the arena.

### Linear track

After finishing open field, mice were water restricted and maintained at a body weight of around 85% of their initial weight. Mice were trained to run back and forth along the track wearing dummy scopes with similar weight as real scopes while experimenters pipetted 10 ml water reward at each end of the track. Mice were habituated to the track for 15min each day for three days. After habituation, imaging session were performed every two days for 3 sessions. Calcium imaging was performed simultaneously with miniCaM imaging of mouse behavior at 50Hz.

### Social interaction

Mice with bilateral relay lens implantations were habituated to the environment and to wearing dummy MiniXL for four days before testing. On the days of test, mice were habituated for 30-60 minutes in their home cage inside the experimental room. During the first session on day 1, a MiniXL was attached to the baseplate and the mouse was placed into an open arena (45cm X 45ch X 30cm). After the subject mouse explored the environment alone for one minute, a novel C57BL/6 male mouse was introduced into the arena as a social target. The two mice interacted freely for seven minutes. The subject mouse was then put back into the homecage alone for five minutes, after which it was transferred back to the same arena. After one minute being alone in the arena, a 3D printed object (10 cm diameter, 10 cm high cone) was placed in the center of the arena as an object target which subject mouse explored for seven minutes. Calcium imaging was performed simultaneously with behavioral recording by a miniCaM both at 22Hz.

### Analysis of behavioral assays Linear track and open field

For experiments involving freely behaving mice, behavioral videos were captured in AVI format using an overhead miniCAM (50 fps). The position of the mouse was extracted by tracking the position of the blue light from the excitation LED. The pixel location of the LED was found by detecting the highest pixel value region within the grayscale behavior recording frames, and then the pixel value location is converted to real-world coordinates.

### Social interaction

Animal’s behavior was recorded by MiniCam from the top of arena at 22Hz. All videos were manually annotated frame by frame to identify the onset and offset of each epoch of social interaction and object exploration. The following criteria must be met for an interaction to be identified as a social or object interaction episode: 1. The subject mouse (mouse wearing miniXL) must initiate investigation of the social target (or object) or respond to the exploration by social target mouse; 2. Social interaction must include approaching, sniffing, grooming, chasing and mounting the social target. The onset of interaction was identified as the frame in which subject nose pointed to the social (object) target within a distance about 1/4-1/3 of its body length. The offset of interaction was identified as the frame in which subject nose started to leave the social (object) target at a distance about 1/4-1/3 of its body length.

### Calcium imaging analysis

Calcium imaging data was analyzed using MiniscopeAnalysis package (https://github.com/etterguillaume/MiniscopeAnalysis) or CalmAn (https://github.com/flatironinstitute/CaImAn-MATLAB) (*27*). The NoRMCorre algorithm was applied to perform motion correction (*28*). The constrained non-negative matrix factorization for microendoscopic data (CNMF-E) approach was used to identify and extract the spatial shapes and fluorescent calcium activity of individual cells(*29*). All cells’ shape and Ca^2+^ traces were manually inspected to ensure high data quality. For imaging data recorded from sessions across days, we applied CellReg package (https://github.com/zivlab/CellReg)^16^ to identify the same cells across days.

### Identifying Place cells Place cells in linear track

For a particular cell to be identified as a place cell in recordings of mice running on the linear track, it must meet the following three criteria: 1. The neuron’s spatial information content must exceed chance levels (P < 0.05) as determined by circularly shuffled distribution analysis; 2. The neuron’s within-session stability (both the first and the second half session) must exceed chance levels (P < 0.05) as determined by circularly shuffled distribution analysis; 3. the spatial activity rate map of the neuron must demonstrate activity in consecutive bins covering a minimum of 10 cm, with activity rates ranking in at least the 95th percentile among the binned activity rates of circular trial shuffled spatial activity rate maps (*13*).

The information content was defined as:

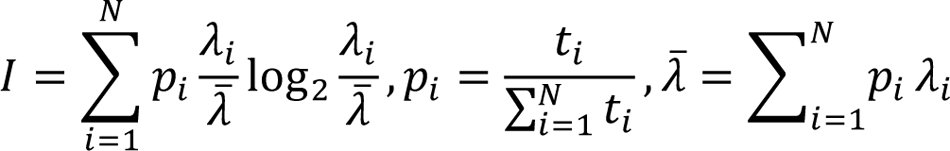

where *t*_*i*_ represents the occupancy time spent in the *i*-th bin and *λ*_*i*_ represents the neural activity rate in the *i*-th bin.

The stability was calculated by taking the Fisher Z-score of the Pearson correlation coefficient between the spatial activity rate maps at two time points (odd versus even trials and first half versus second half of the trials).

Spatial neural activity rates were calculated using 2-cm-wide spatial bins and a speed threshold of greater than 10 cm s−1. Temporal neural activity and occupancy of the animal were spatially binned and then smoothed using a Gaussian kernel with σ = 5 cm. The binned neural activity was divided by the binned occupancy to calculate the spatial neural activity rate of each cell.

### Place cells in open field arena

To identify place cells in mice exploring a 2-dimensional environment, the following criteria needed to be met: 1. The spatial information content need to be significantly above chance (P < 0.05) based on the circularly shuffled distribution; the neuron’s within-session stability both the first and the second half session) must have exceeded chance levels (P < 0.05); and the spatial activity rate map of the neuron must have covered five adjacent bins (*14*).

### Identifying social cells/object cells

Imaging data and behavior videos were synchronized using timestamps derived from the computer’s internal clock. For each cell, calcium traces were normalized to the maximum value in the entire recording session. We used receiver operating characteristic (ROC) analysis (*30*, *31*) to identify social interaction responsive cells and object exploration responsive cells. For each cell, we calculated an area under ROC curve (auROC) value (0–1) to measure the overlap of calcium traces with binarized social and object interaction plots and compared these values to those obtained from 1000 times randomly shuffled traces. Neurons with auROC values over 97.5 percentile of shuffled data were classified as social excited cells (SE) or object excited cells (OE). Neurons with auROC values below the 2.5 percentile of shuffle data were classified as social inhibited cells (SI) or object inhibited cells (OI). The remaining neurons were classified as non-modulated neurons (SN, ON).

### Statistics

All statistical analyses were conducted using GraphPad Prism 9 and MATLAB (R2020a). All plots were presented in mean ± SEM unless otherwise specified. Statistical significance was defined with P<0.05. All statistical methods and sample numbers are described in individual figure legends.

## Author contributions

D.A. and C.G. conceived the project. D.A. and C.G. developed MiniXL. P.Z. performed all experiments. C.G. assisted performing experiments. P.Z. and C.G. designed the study, analyzed data and wrote manuscript. M.X. redesigned the PCB circuit, updated the software, and did the fundamental test of the device. L.C. assisted in the figures and manuscript. P.G. and D.A. supervised the entire project.

## Supplemental Materials

**Fig. S1.**
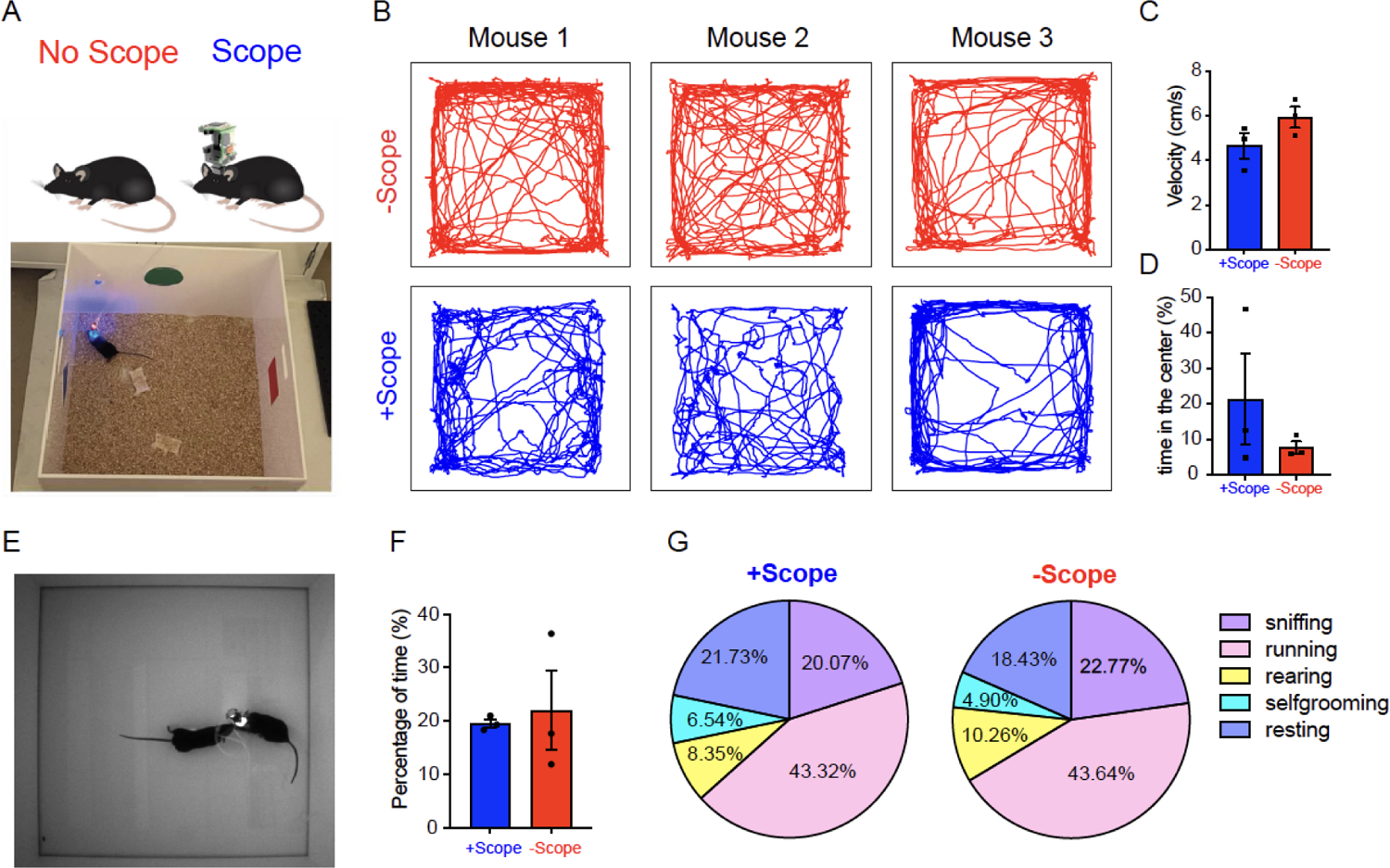
MiniXL does not limit mouse natural behavior. (A) Schematic diagram of mouse inside open field area with visual cues. (B) Trajectories of three mice with or without wearing MiniXL. (C) Velocity of mice travelling in the arena with or without wearing MiniXL. (D) Percentage of time mice spent in the center area of the chamber with or without MiniXL. (E) Schematic diagram of mice social interaction test. (F) Percentage of time subject mice interacting with social target with or without MiniXL. (G) Pie chart showed percentage of different types of natural behavior subject mice performed. N=3 animals.

**Fig. S2.**
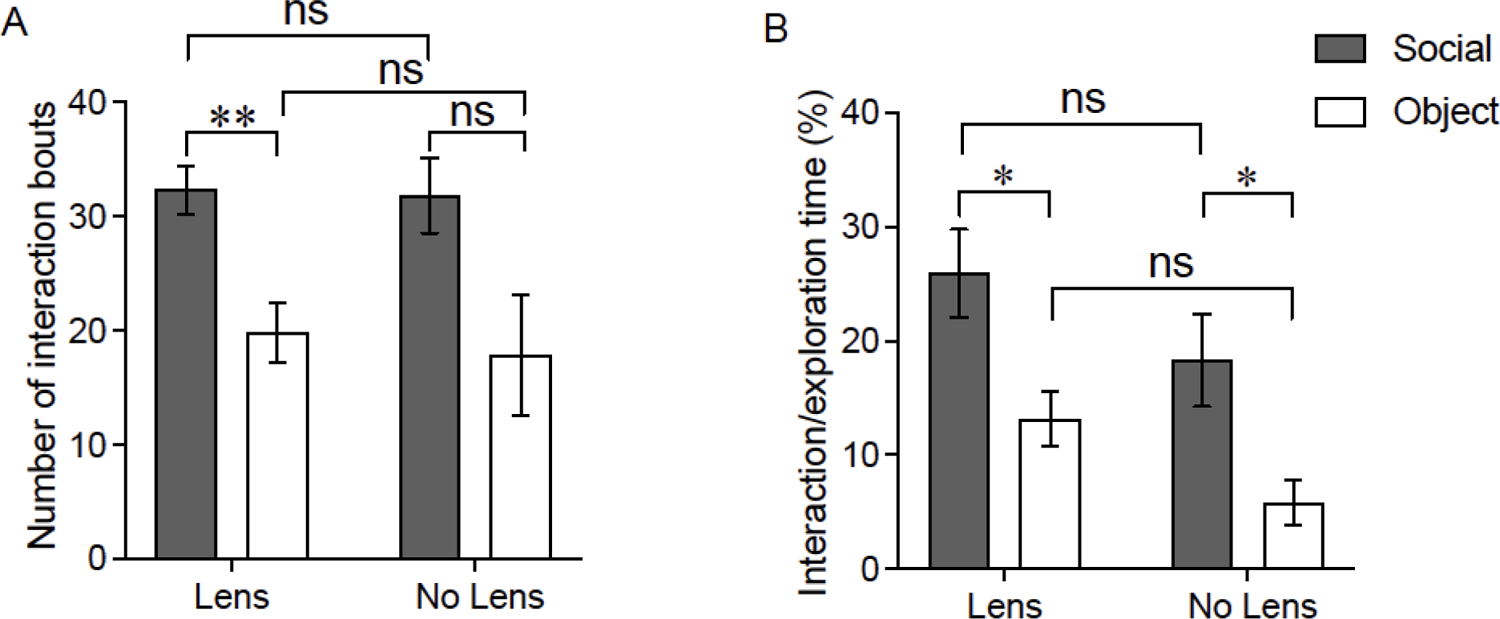
Lens implantation does not change mice social behavior. (A) Number of interaction bouts initiated by subject mice to social target and object. (B) Percentage of time mice spent on social interaction and object exploration. With lens implantation, N=11, Without lens implantation, N=6. * P<0.05, ** P<0.05. Mice had lens implanted into NAc and wearing V4 miniscope.

**Fig. S3.**
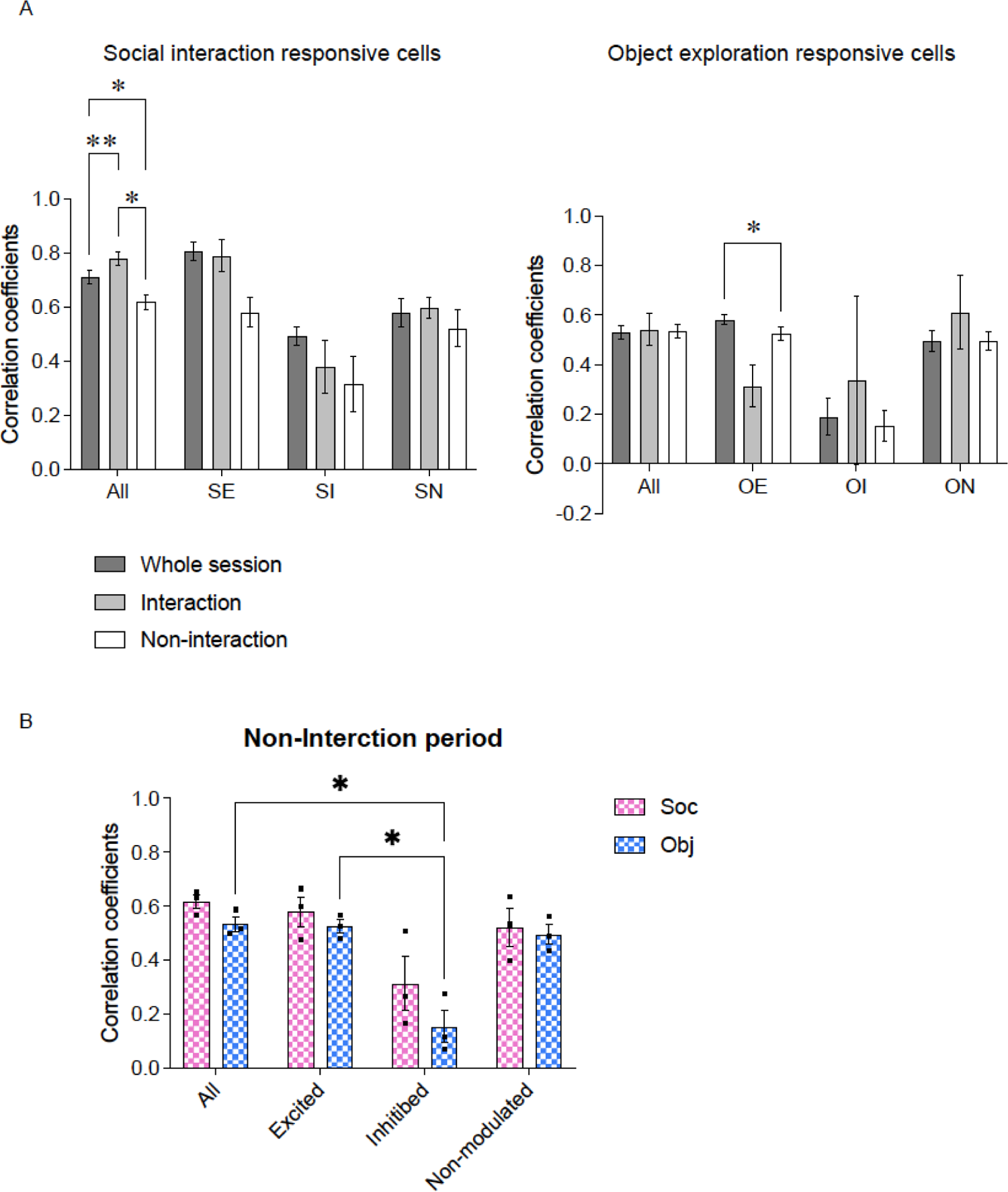
Correlation coefficients between left and righ mPFC neurons. (A) Correlation coefficients of left and right mPFC neurons during social interaction sessions (left panel) and object exploration session (right panel). (B) Correlation coefficients of left and right mPFC neurons during non-interaction period of social session and object session (white bars in A).

**Fig. S4.**
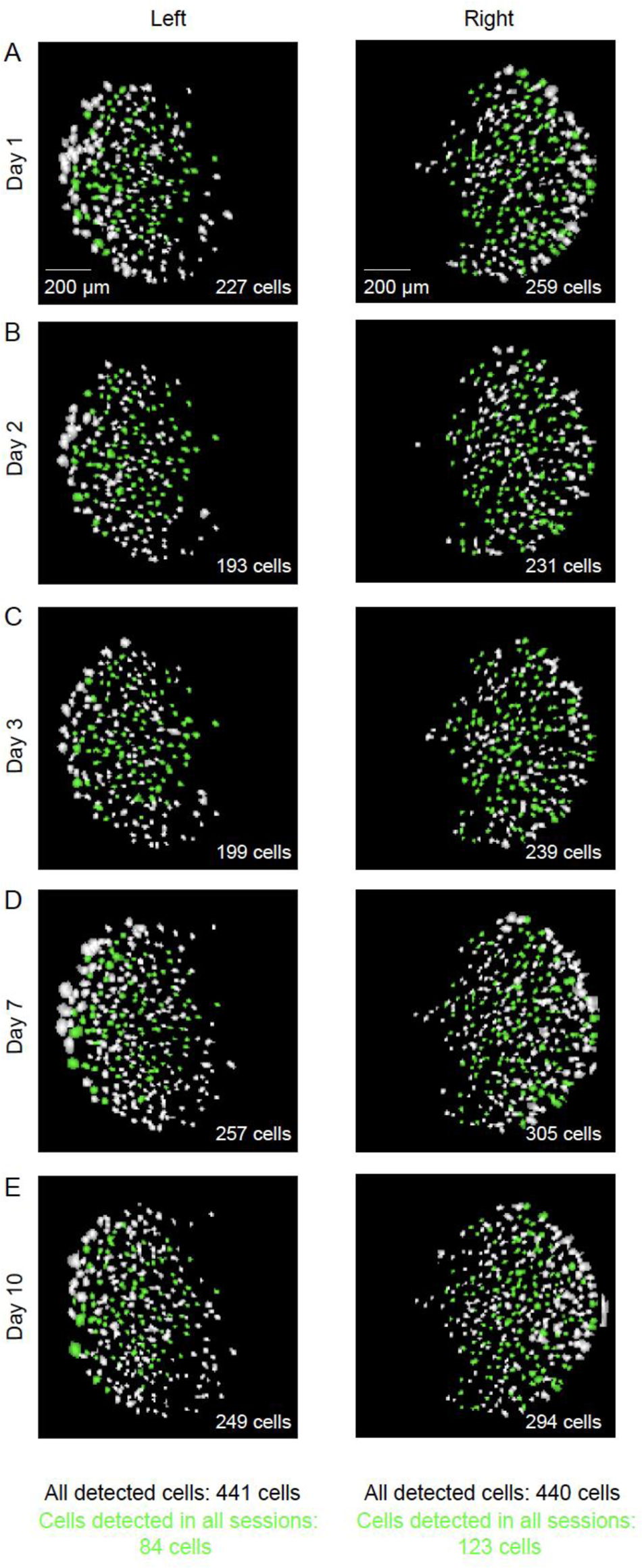
Bilateral imaging of PFC in freely social interacting mice across multiple days using MiniXL. A-E CNMF-E extracted cells from left and right mPFC on different days. Green labelled cells are those detected from all sessions.

---

**Movie S1.** Raw video of Calcium imaging recorded from dCA1 when mouse running on the linear track.

**Movie S2.** Raw video of Calcium imaging recorded from bilateral mPFCs during mouse social interaction test.

**Movie S3.** Calcium imaging recorded from NAc and PFC simultaneously synchronized with animal behavior during open field test. Upper left: screen shot of cropped field of view (left, NAc; right, mPFC); Upper middle and Right: motion corrected videos of NAc and mPFC calcium signals; Bottom: Trajectory of the mouse running in the chamber.

